# The elevational ascent and spread of invasive annual grass dominance in the Great Basin, USA

**DOI:** 10.1101/2021.01.05.425458

**Authors:** Joseph T. Smith, Brady W. Allred, Chad S. Boyd, Kirk W. Davies, Matthew O. Jones, Andrew R. Kleinhesselink, Jeremy D. Maestas, Scott L. Morford, David E. Naugle

## Abstract

**Aim:** In the western US, sagebrush (*Artemisia* spp.) and salt desert shrublands are rapidly transitioning to communities dominated by exotic annual grasses, a novel and often self-reinforcing state that threatens the economic sustainability and conservation value of rangelands. Climate change is predicted to directly and indirectly favor annual grasses, potentially pushing transitions to annual grass dominance into higher elevations and north-facing aspects. We sought to quantify the expansion of annual grass-dominated vegetation communities along topographic gradients over the past several decades.

**Location:** Our analysis focused on rangelands among three ecoregions in the Great Basin of the western US, where several species of exotic annual grasses are widespread among shrub and perennial grass-dominated vegetation communities.

**Methods:** We used recently developed remote sensing-based rangeland vegetation data to produce yearly maps of annual grass-dominated vegetation communities spanning the period 1990–2020. With these maps, we quantified the rate of spread and characterized changes in the topographic distribution (i.e., elevation and aspect) of areas transitioning to annual grass dominance.

**Results:** We documented more than an eight-fold increase in annual grass-dominated area (to >77,000 km^2^) occurring at an average rate of >2,300 km^2^ yr^-1^. In 2020, annual grasses dominated one fifth (19.8%) of Great Basin rangelands. This rapid expansion is associated with a broadening of the topographic niche, with widespread movement into higher elevations and north-facing aspects.

**Main conclusions:** Accelerated, strategic intervention is critically needed to conserve the fragile band of rangelands being compressed between annual grassland transitions at lower elevations and woodland expansion at higher elevations.

## Introduction

Grasses are highly successful invaders globally with the capacity to dramatically reshape rangelands (D’Antonio & Vitousek, 1992; Godfree et al., 2017). Among diverse ecosystems across several continents, consequences of grass invasions include increased risk to human life and property from larger and/or more frequent wildfires (Fusco et al., 2019), impacts on human health (Johnston et al., 2009), disruption of hydrologic and nutrient cycles (Evans et al., 2001; Germino et al., 2016; Rossiter-Rachor et al., 2009), loss of habitat for sensitive species (Coates et al., 2016), and reduced biodiversity across trophic levels (K. W. Davies, 2011; Pyšek et al., 2012). Commonly, disruptions to invaded communities are sufficient to force transitions into alternative stable states characterized by dominance of exotic grasses (e.g., “grass-fire cycles;” D’Antonio & Vitousek, 1992). Despite advancements in our understanding of the mechanisms and outcomes of exotic grass invasions on recipient ecosystems, sporadic monitoring has limited our ability to quantify the scope and pace of such ecosystem transformations at broad spatio-temporal scales (Bromberg et al., 2011; Levick et al., 2015).

In the arid and semi-arid Great Basin of the western U.S., exotic annual grasses including cheatgrass (*Bromus tectorum*), red brome (*B. rubens*), medusahead (*Taeniatherum caput-medusae*), and ventenata (*Ventenata dubia*) have become virtually ubiquitous (Mack, 1981; Nicolli et al., 2020; Young & Evans, 1970). These grasses colonize interstices between native perennial bunchgrasses and shrubs, increasing the amount and continuity of fine fuels (K. W. Davies & Nafus, 2013). Consequently, annual grass-invaded vegetation communities burn 2–4 times more frequently than uninvaded communities (Balch et al., 2013; Bradley et al., 2018). Post-fire reestablishment of native vegetation often proves exceedingly challenging due in part to pre-emptive resource use by early-germinating annual grasses (K. W. Davies, 2010; Eliason & Allen, 1997; Melgoza et al., 1990). Ultimately, this cycle of invasion, fire, and exclusion of native competitors can push recipient communities across a threshold into an undesirable state of dominance by exotic annual grasses (hereafter, annual grass dominance). Similar to exotic grass invasions globally (Godfree et al., 2017), transition to annual grass dominance erodes both the economic and conservation value of Great Basin landscapes (Knapp, 1996).

Climate change may facilitate annual grass dominance in the Great Basin. Warmer temperatures and earlier snowmelt are predicted to favor establishment, growth, and reproduction of annual grasses throughout much of the region (Blumenthal et al., 2016; Bradley, 2009; Compagnoni & Adler, 2014b). Larger and more frequent wildfires resulting from extended fire seasons may accelerate transitions to annual grass dominance (Abatzoglou & Kolden, 2011). Potentially exacerbating these dynamics, rising atmospheric CO_2_ may increase annual grass biomass and flammability (Ziska et al., 2005). The net effects of climate change on the distribution of annual grass dominance are uncertain, however, as effects of warming are species-specific (Bradley et al., 2016; Bykova & Sage, 2012), contingent upon soil moisture (Compagnoni & Adler, 2014a; Thill et al., 1979; Zelikova et al., 2013), and moderated by the response of the recipient native vegetation community (Chambers et al., 2007; Compagnoni & Adler, 2014b). Range contractions of certain species are predicted under some climate scenarios (Bradley, 2009), but vacated regions may subsequently be invaded by other exotic annual grasses (Bradley et al., 2016; Bykova & Sage, 2012).

Elevational range shifts are among hypothesized responses of annual grasses to climate change (Bradley et al., 2016; Compagnoni & Adler, 2014a, 2014b). The Great Basin is mountainous, with elevations ranging from <700 m to >3000 m. Historically, physiological constraints have limited the spread of *B. tectorum* at higher elevations and on cooler, north-facing slopes (Chambers et al., 2007, 2014). As temperatures warm, timing of snowmelt advances, and precipitation increasingly falls as rain, this may lead higher elevations to become suitable for establishment and growth of *B. tectorum* (Compagnoni & Adler, 2014a; Concilio et al., 2013). Furthermore, warming and increasing summer aridity has increased fire in higher elevations across mountainous ecosystems of the western US (Alizadeh et al., 2021), a phenomenon which could indirectly assist the spread of annual grasses into higher elevations.

The spatio-temporal dynamics of state transitions to annual grass dominance in the Great Basin remain poorly understood. Remote sensing has been successfully used to map annual grass dominance at ecoregional extents (Bradley & Mustard, 2005; Peterson, 2005), but these occasional snapshots in time provide limited insight into the rate or trajectory of transitions along bioclimatic gradients. An understanding of these dynamics is needed to inform management and policy. Using a recently developed remotely sensed vegetation cover product that provides continuous coverage of western US rangelands across space and time (Allred et al., 2020; Jones et al., 2020), we quantify the expansion of annual grass dominance in the Great Basin and test for shifts in distribution along topographic gradients over the past several decades.

## Methods

### Study area

We characterized expansion of annual grass dominance within three level III ecoregions (Omernik & Griffith, 2014) that overlap the hydrologic Great Basin and share similar climates and potential vegetation (Fig. 1). These include the Central Basin and Range (CBR), Northern Basin and Range (NBR), and Snake River Plain (SRP). Within these ecoregions, we limited our analyses to rangeland land cover as defined by Reeves and Mitchell (2011). We also excluded agricultural land cover identified as hay, alfalfa, and idle cropland in the Cropland Data Layer (USDA National Agricultural Statistics Service, 2020).

**Figure 1.**
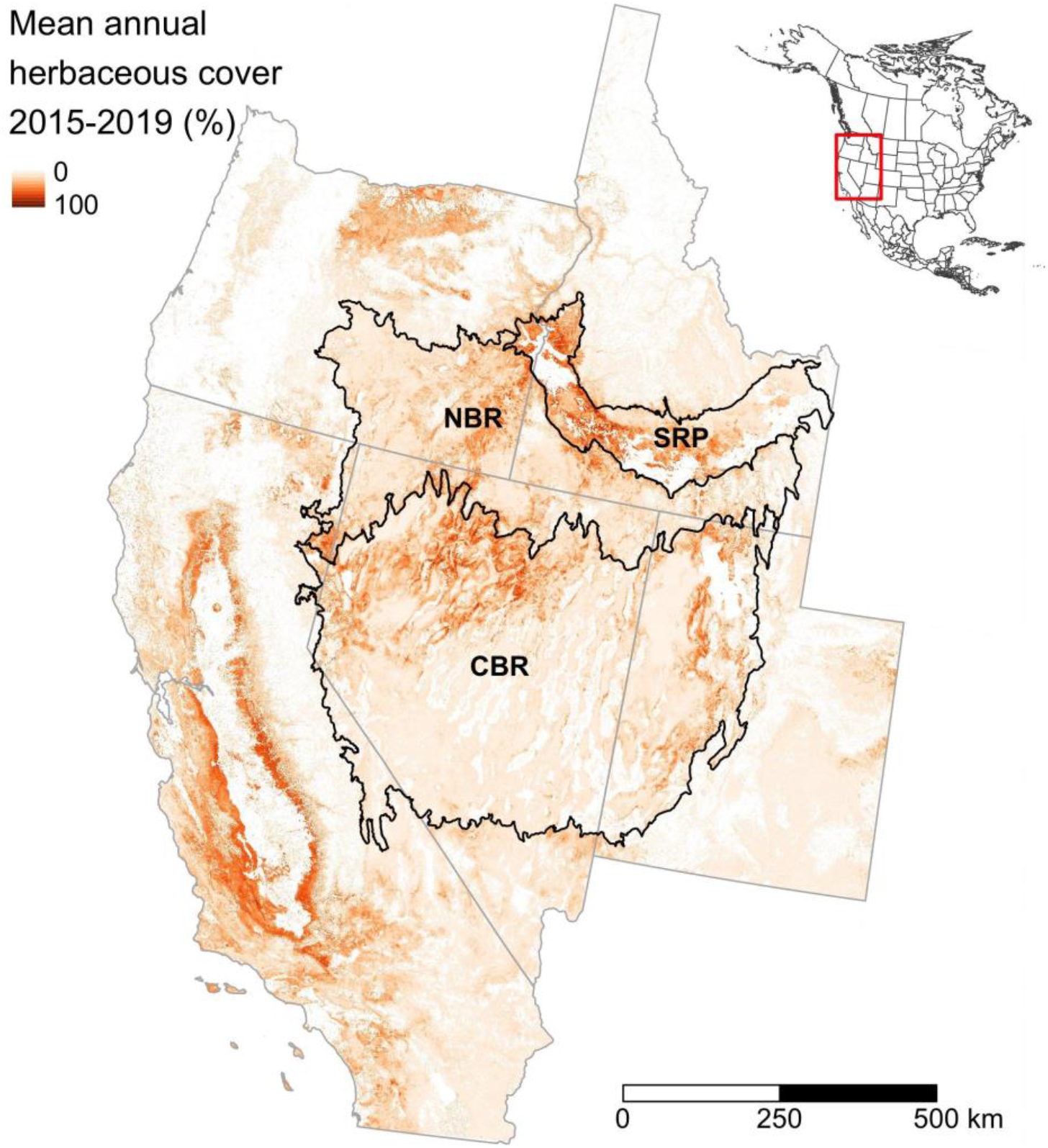
The Great Basin covers >500,000 km^2^ and encompases parts of 5 US states. Our analysis included the Central Basin and Range (CBR), Northern Basin and Range (NBR), and Snake River Plain (SRP) ecoregions.

### Classification of invasive annual grass-dominated communities

We used a remote sensing product, the Rangeland Analysis Platform (RAP; Allred et al., 2021; Jones et al., 2018) to produce high resolution (30 m) maps of annual grass dominance throughout the Great Basin for each year from 1984-2019. RAP uses Landsat satellite imagery to estimate areal canopy cover (hereafter cover) of major plant functional groups, specifically, perennial forbs and grasses (PFGC), annual forbs and grasses (AFGC), shrubs (SHR), bare ground (BG), litter (LTR), and trees (TREE).

Cover of herbaceous vegetation in arid environments changes rapidly both within and between years, potentially confounding efforts to quantify trends and identify state transitions (Bradley & Mustard, 2005). To focus our analysis on long-term changes in vegetation state, we used a smoothing algorithm to smooth the time series of cover estimates for each pixel. We used Holt’s linear exponential smoothing method (Holt, 2004) and selected smoothing parameters by applying them to 1,000 randomly selected pixels from within the study area and selecting the parameters that minimized mean absolute error between one-step ahead cover estimates produced by the smoothing function and raw cover values. This dampened sharp year-to-year fluctuations while still revealing longer-term trends and abrupt shifts following disturbances (e.g., wildfire; Fig. S1).

Although vegetation may fall anywhere along a continuum from uninvaded to complete annual grass dominance, we sought to identify highly-invaded communities where herbaceous vegetation is predominantly (i.e., >50% by cover) exotic annual grasses and cover of woody vegetation is absent or diminished. This severely invaded state is thought to represent an equilibrium from which a return back to a perennial-dominated state with some shrub cover is unlikely without intensive management (S. Bagchi et al., 2013; G. M. Davies et al., 2012). We took a data-driven approach to identifying which pixels fell into this annual grass dominated state following the analytical framework outlined by Bagchi et al. (2013; 2012). We used *k*-means++ clustering (Arthur & Vassilvitskii, 2007) to group pixels into a finite number of clusters representing discrete vegetation ‘states’, and selected the number of clusters (*k*) using the silhouette method (Rousseeuw, 1987). We then examined the characteristics of the resulting clusters to identify the cluster most closely associated with annual grass dominance. Selection of *k* and characterization of clusters was performed in R on 10,000 pixels sampled randomly in time and space from the raw RAP cover time series. Once we determined *k*, training and clustering of the smoothed RAP cover time series and all other raster processing steps were completed in Google Earth Engine (GEE; Gorelick et al., 2017) via the GEE code editor. We clustered the RAP cover time series beginning in 1990, such that cover estimates from 1984– 1989 served to initialize smoothing.

### Validation

We validated our annual grass dominance mapping procedure by comparing the remote sensing-derived vegetation cluster dataset to an independent (i.e., not used in the training of RAP models) set of 1,486 field-measured monitoring plots from the US Bureau of Land Management Assessment, Inventory, and Monitoring program (AIM; Taylor et al., 2014). We applied the same trained clusterer used on the smoothed RAP cover time series to the field data to determine the cluster identity of each of the field plots. From these cluster labels, we assigned each plot a binary class label distinguishing annual grass dominance from other clusters. The resulting field plot-based classes (hereafter field class) were then compared to the smoothed RAP imagery-based classes (hereafter RAP class) from the year corresponding to the sampling date. Agreement between plot classes and RAP classes was assessed with statistics based on the resulting confusion matrix, including kappa, sensitivity/specificity and positive/negative predictive value.

### Characterizing trends

We summed the area of pixels in the annual grass-dominated cluster in each year to derive yearly estimates of the areal extent of annual grass dominance within each ecoregion. To quantify trends in extent of annual grass dominance, we used the non-parametric Theil–Sen estimator (Sen, 1968; Theil, 1950). Estimates and 95% confidence intervals of the Theil–Sen estimator were calculated using the `sens.slope` function in the ‘trend’ package (Pohlert, 2020) in R.

We examined trends in elevation and aspect to test whether the topographic distribution of pixels transitioning to annual grasslands changed over the duration of the time series. For pixels classified as annual grassland in ≥1 yr, we determined the earliest year in the time series that the pixel was assigned to the annual grassland cluster (transition year). Omitting pixels already classified as annual grasslands at the beginning of the time series (1990), we randomly sampled *n* = 500 pixels from each transition year in each ecoregion and determined their elevation and aspect from the USGS National Elevation Dataset (NED) ⅓ arc-second digital elevation model. We used the cosine of aspect (in radians, θ) as an index, ranging from -1 to 1, differentiating north-facing slopes (cos(θ) > 0) from south-facing slopes (cos(θ) < 0). Hereafter we refer to cos(θ) as ‘northness.’ To characterize trends in elevation, we used quantile regression to estimate temporal trends in the 10th (*τ* = 0.1), 50th (*τ* = 0.5), and 90th percentiles (*τ* = 0.9) of elevations. We used AIC to select from among several *a priori* models of the relationship between elevation, northness, year, and ecoregion, including two-and three-way interactions (candidate models and AIC provided in Table S1 in Supporting Information). We estimated regression coefficients with the ‘quantreg’ package (v 5.67; Koenker et al., 2020) in R, and computed standard errors and confidence intervals from 1,000 bootstrap samples.

Masking out slopes < 5°, we calculated the empirical cumulative distribution function (ECDF) of the aspects of all pixels transitioning to annual grass dominance during each decade spanned by the time series (i.e., 1991–2000, 2001–2010, and 2011–2020). We display these decadal ECDFs grouped by elevation band (terciles of rangelands assigned to the annual grass dominance cluster in ≥1 yr) and ecoregion.

## Results

### Classification

Using smoothing parameters *α* = 0.25 and *β*\* = 0.01 for Holt’s smoothing method minimized MAE across the 1,000 tested AFGC time series (see Fig. S1 for examples of smoothing). Average MAE (i.e., the average absolute difference between the smoothed value and raw RAP estimate for a pixel-year) was 4.7%.

Using *k*-means++ clustering and the silhouette method, we identified *k* = 4 as the optimal number of clusters in the data (Fig. S2). Cluster 0 was characterized by high annual forb and grass cover (median AFGC = 29%), annual dominance (median ratio AFGC:PFGC = 1.57:1), and low shrub cover (median SHR = 9%), therefore this cluster was selected to represent annual grass dominance (Fig. 2, Table 1). The other 3 clusters identified areas mostly devoid of vegetation (cluster 3), areas composed predominantly of bare ground and shrubs (cluster 2), and areas characterized by varying degrees of co-dominance of shrubs and perennial grasses and forbs (cluster 1; Table 1). Although clusters occupied distinct regions of the multivariate space, boundaries between clusters were more-or-less continuous (Fig. 2). Pixels assigned to cluster 1 often displayed substantial cover of annual forbs and grasses (median = 7%), but they consistently comprised the minor component of herbaceous vegetation cover (median ratio AFGC:PFGC = 0.2:1, 95th percentile = 0.76:1).

**Figure 2.**
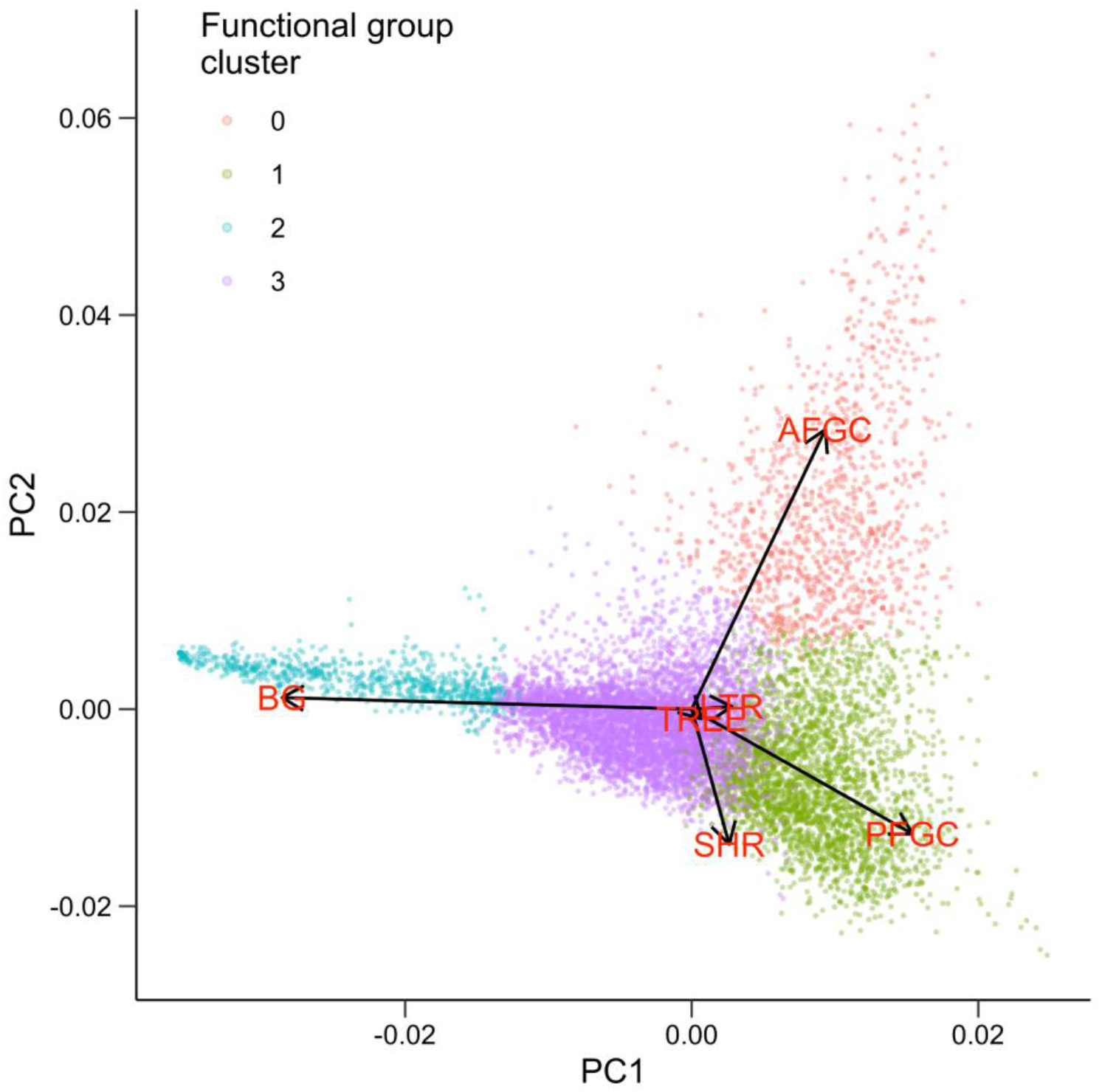
Composition of *k* = 4 functional group cover clusters along the first two principal components, which explained 77% of the total variance. Vectors indicate loadings of all function group cover components (AFGC = annual forb and grass cover; BG = bare ground; LTR = litter; PFGC = perennial forb and grass cover; SHR = shrub cover; TREE = tree cover).

**Table 1.** Composition (percent cover) of Great Basin rangeland functional group clusters identified via *k*-means partitioning. Percentiles were calculated from *n* = 10,000 randomly selected pixels.

| Cluster ID | Functional group<sup>a</sup> | Percentile |  |  |  |  |
| --- | --- | --- | --- | --- | --- | --- |
|  |  | 5 | 25 | 50 | 75 | 95 |
| 0<sup>b</sup> | AFGC | 12 | 21 | 29 | 38 | 57 |
|  | BG | 3 | 7 | 11 | 16 | 27 |
|  | LTR | 8 | 12 | 15 | 18 | 26 |
|  | PFGC | 4 | 11 | 18 | 27 | 40 |
|  | SHR | 3 | 6 | 9 | 15 | 23 |
|  | TREE | 0 | 0 | 0 | 1 | 3 |
| 1 | AFGC | 0 | 3 | 7 | 12 | 22 |
|  | BG | 2 | 5 | 9 | 14 | 24 |
|  | LTR | 4 | 9 | 12 | 15 | 19 |
|  | PFGC | 17 | 25 | 32 | 44 | 73 |
|  | SHR | 2 | 11 | 18 | 26 | 40 |
|  | TREE | 0 | 0 | 1 | 2 | 16 |
| 2 | AFGC | 0 | 3 | 5 | 10 | 21 |
|  | BG | 7 | 15 | 23 | 31 | 41 |
|  | LTR | 7 | 11 | 13 | 15 | 18 |
|  | PFGC | 1 | 4 | 8 | 12 | 19 |
|  | SHR | 6 | 13 | 18 | 24 | 32 |
|  | TREE | 0 | 0 | 0 | 4 | 38 |
| 3 | AFGC | 0 | 0 | 1 | 2 | 5 |
|  | BG | 46 | 57 | 74 | 87 | 100 |
|  | LTR | 0 | 0 | 4 | 6 | 11 |
|  | PFGC | 0 | 2 | 4 | 6 | 16 |
|  | SHR | 0 | 0 | 5 | 9 | 14 |
|  | TREE | 0 | 0 | 0 | 0 | 1 |

<sup>a</sup> AFGC = annual forb and grass cover; BG = bare ground; LTR = litter; PFGC = perennial forb and grass cover; SHR = shrub cover; TREE = tree cover.
<sup>b</sup> Cluster selected to represent annual grasslands.

### Validation

Functional group composition of field plots assigned to the annual grass dominance cluster are shown in Fig. S2. Agreement between field and RAP binary class assignment (annual grass-dominated vs other) was good. Using RAP class as the prediction and field class as the reference, accuracy (percent correctly classified) was 0.88, and kappa was 0.66. Sensitivity, or the fraction of true positive cases (i.e., annual grass dominance according to field data) correctly predicted to belong to the annual grass dominance cluster, was 0.64. Specificity, or the fraction of true negative cases (not annual grass dominance according to field data) correctly assigned to non-annual grass dominance categories, was 0.97. Positive predictive value, or the fraction of predicted positive cases that were actually annual grass-dominated, was 0.88, and negative predictive value was 0.88. The substantially lower sensitivity relative to specificity suggests the true extent of annual grass dominance at any point in time is underestimated by our methods.

This discrepancy was partly a consequence of smoothing prior to clustering. The smoothed value in a given year is a weighted function of that year’s estimate and past years’ estimates, with the weight decaying into the past. When cover of a functional group trends quickly in either direction, the smoothed value will lag behind in reflecting that change (Fig. S1). Because the extent of annual grass dominance is generally increasing and sometimes increases rapidly following disturbance, smoothed annual forb and grass cover values will tend to underestimate current cover and the transition of pixels into the annual grass dominance cluster will lag behind the true transition on the ground by a short period.

To confirm this, we compared the field classes to RAP classes 1 yr after field measurements were made. As expected, agreement between RAP and field class assignments improved: kappa increased to 0.69 and sensitivity increased to 0.68, with little change in specificity (0.96). Still, a false negative rate of >0.3 indicates a systematic underestimation of the extent of annual forb and grass-dominated vegetation communities.

### Trends

The area of annual grass dominance in the Great Basin increased >8-fold during the study period, from 8,964 km^2^ in 1990 to 77,617 km^2^ in 2020 (Fig. 3). Aggregated across the Great Basin, this represents an estimated annual rate of increase of 2,373 km^2^ yr^-1^ (95% CI = 2,055–2,638 km^2^ yr^-1^). At the ecoregional scale, estimated annual rates of increase ranged from 430.2 km^2^ yr^-1^ (95% CI = 370.1–485.8 km^2^ yr^-1^) in SRP to 1,202.8 km^2^ yr^-1^ (95% CI = 1,015.5– 1,368.5 km^2^ yr^-1^) in CBR (Fig. 3). In 2020, the annual grass dominance cluster occupied 17% (41,537 km^2^) of rangelands in CBR, 18% (21,771 km^2^) of rangelands in NBR, and 43% (14,309 km^2^) of rangelands in SRP, or 19.8% of rangelands across ecoregions.

**Figure 3.**
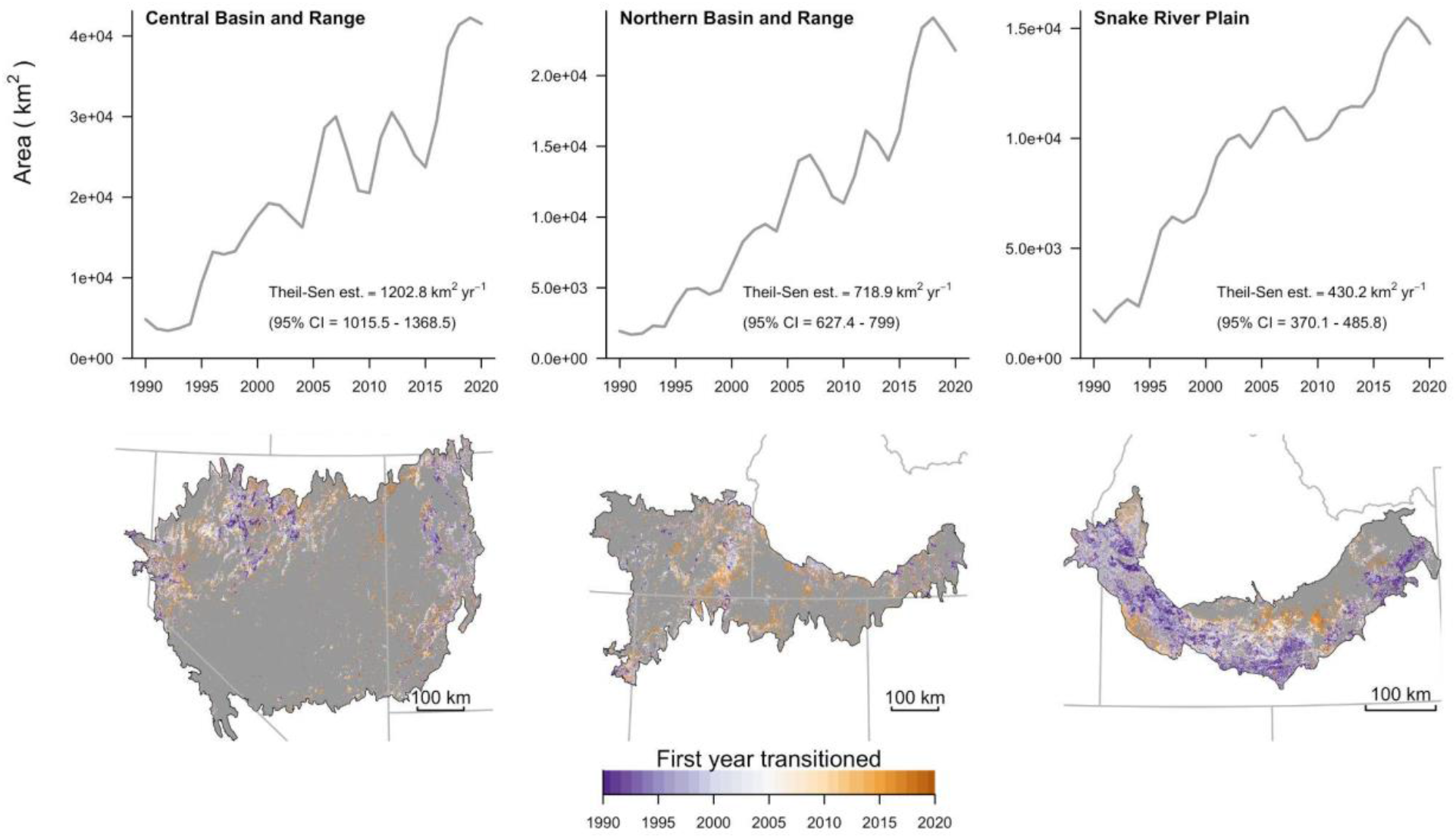
Growth in area of invasive annual grass-dominated vegetation communities in the Great Basin, USA, 1990–2020 (top row). Spatial distribution of pixels characterized by annual grass dominance in ≥1 yr, with colors indicating the first year transitioned (bottom row). Darkest purple indicates transition in or prior to 1990, the first year in the time series.

Elevations of transitions to annual grass dominance increased through time (Table 2; Fig. 4), though the rate of increase differed by ecoregion and aspect (Table 3). For all quantiles, the three-way year × northness × ecoregion interaction model received unequivocal support based on AIC (Table S1). Elevations of transitions were highest in the most southerly ecoregion (CBR) and decreased through the most northerly ecoregion (SRP). On east- and west-facing aspects (i.e., setting northness = 0), the median elevation of transitions (*τ* = 0.5) increased 4.7 m yr^-1^ (95% CI = 4.2–5.2 m yr^-1^) in CBR, 6.6 m yr^-1^ (95% CI = 6.1–7.2 m yr^-1^) in NBR, and 14.8 m yr^-1^ (95% CI = 14.1–15.4 m yr^-1^) in SRP.

**Table 2.**
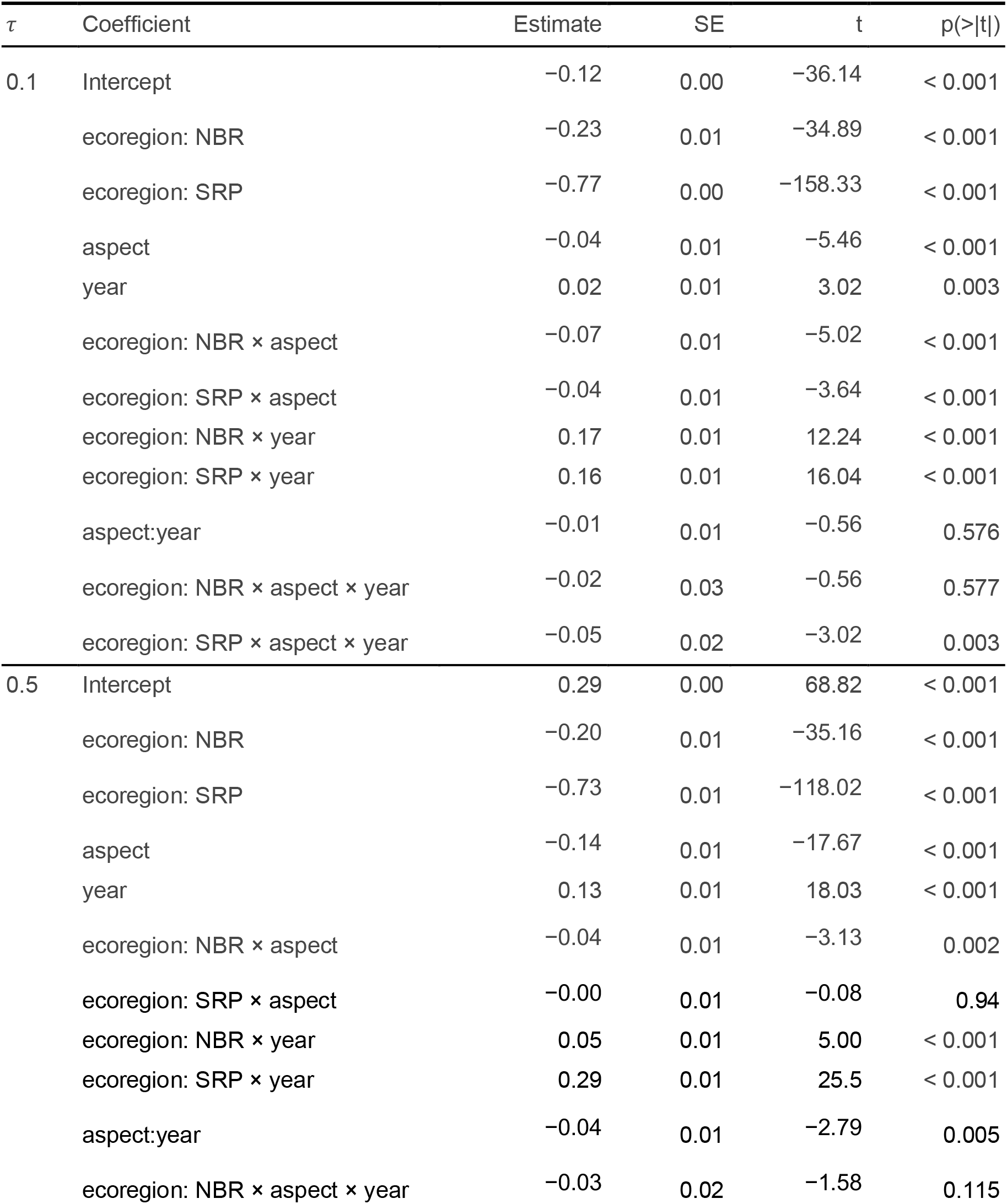
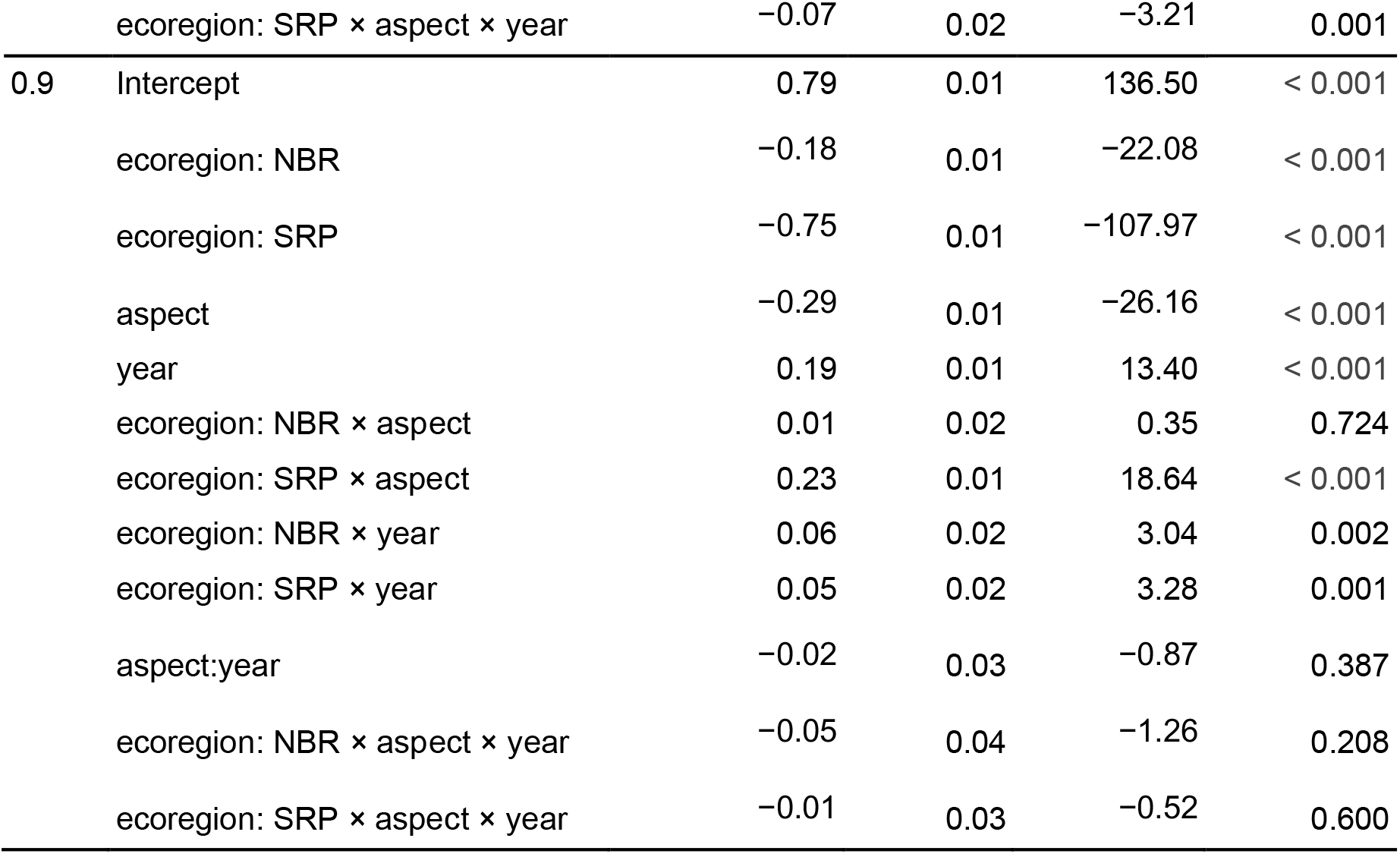
Coefficient estimates of quantile regression models describing relationships between elevation, aspect, and year of transition to invasive annual grass dominance among three Great Basin ecoregions, 1990–2020. Continuous variables were standardized by subtracting the mean and dividing by two standard deviations.

**Figure 4.**
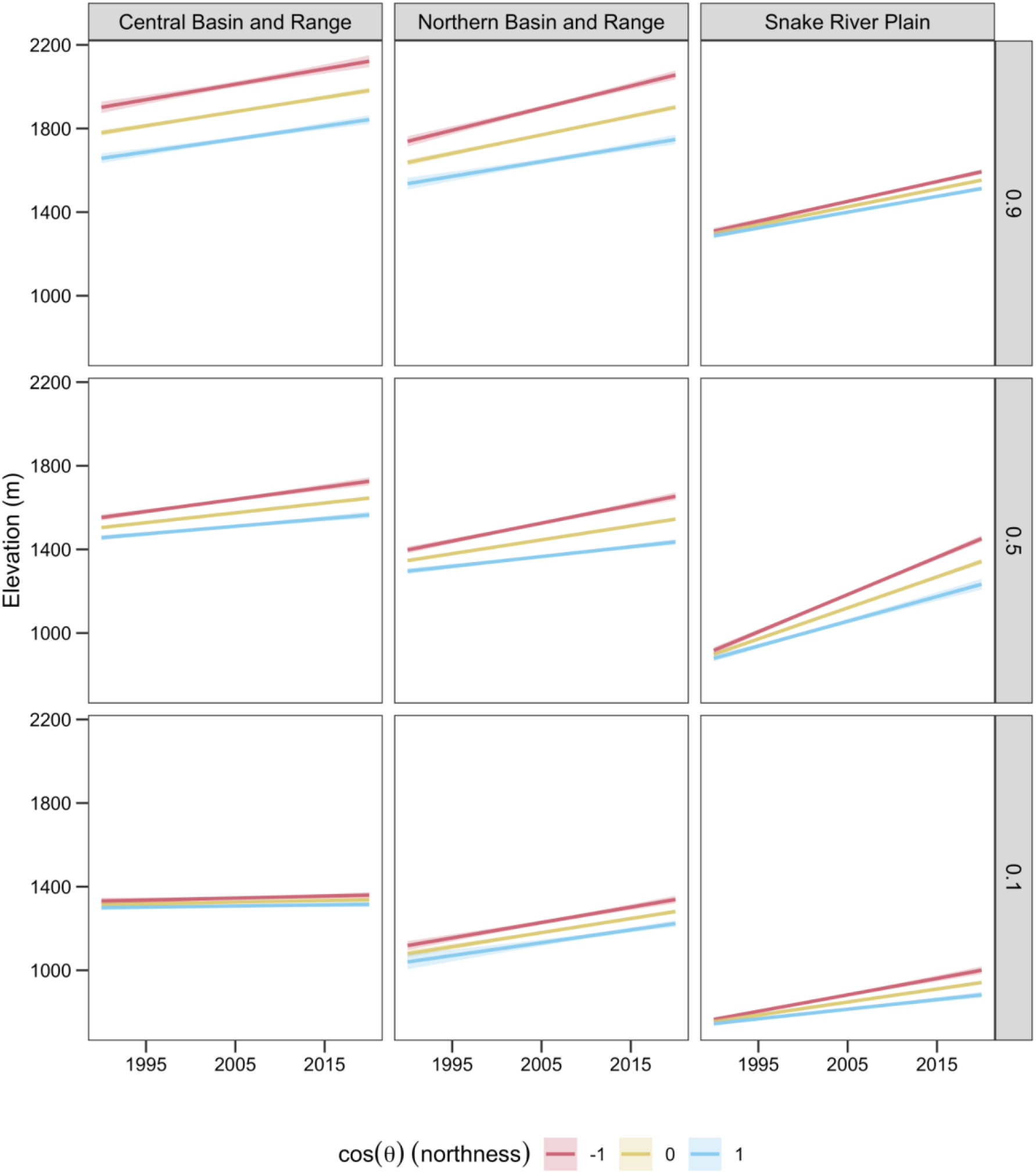
Trends in the 0.1 (top row), 0.5 (middle row), and 0.9 (bottom row) quantiles of elevation of transitions to exotic annual grass dominance in the Great Basin, USA, 1990–2020. Trends in all quantiles were best explained by a three-way interaction between year, aspect, and ecoregion.

Empirical CDFs revealed substantial shifts toward more north-facing aspects through time across elevation bands and ecoregions, with the largest shifts during the last decade (Fig. 4). The smallest aspect shifts occurred at low and mid elevations in CBR, and the largest shifts were seen across elevations in NBR, at mid elevations in SRP, and at high elevations in CBR (Fig. 4).

## Discussion

Annual grass dominance has expanded with alarming speed in recent decades, increasing eight-fold in area since 1990 in the Great Basin. We estimate annual grass dominance now characterizes one fifth (19.8%, >77,000 km^2^) of Great Basin rangelands, and has expanded by >2,300 km^2^ annually, a rate proportionally greater than recent deforestation of the Amazon^2^. The most rapid growth occurred in the last decade (2011-2020), averaging >3,700 km^2^ annually across the Great Basin. Consistent with predictions based on warming trends, movement into higher elevations has allowed expansion to continue more-or-less unabated (Fig. 3). This steady ascent of annual grasses now threatens higher elevation rangelands formerly thought to be minimally vulnerable to transition (Chambers et al., 2007; Johnson et al., 2019).

Our estimates of the extent of annual grass dominance were broadly consistent with previous studies, though differences among study areas make direct comparisons imprecise. For example, Pellant & Hall (1994) estimated 10,027 km^2^ (2,477,837 ac) of Bureau of Land Management rangelands across Idaho, Oregon, Utah, and Nevada was dominated by either *B. tectorum, T. caput-medusae*, or a combination of the two species, in the early 1990s (table 2). Our estimate identifies 8,964 km^2^ of annual grass dominance across all land ownerships within the Great Basin in 1990. Bradley & Mustard (2005) estimated 20,000 km^2^ was dominated by *B. tectorum* across the hydrologic Great Basin, a region roughly corresponding to the Central and Northern Basin and Range ecoregions, using a time series of Landsat imagery through 2001. Our method places the extent of annual grass dominance in those two ecoregions at 27,500 km^2^ in 2001.

Elevational ascent of annual grass dominance has long been assumed (e.g., Concilio & Loik, 2013; K. W. Davies et al., 2011), but our analysis is the first to empirically confirm this widespread phenomenon by capitalizing on advanced remote sensing products. Across the Great Basin, the median elevation of transitions to annual grass dominance has increased by 198–445 m (650–1,460 ft) over the last 30 years (Fig. 4). Ascent was faster on hot and dry south-facing aspects historically most susceptible to invasion (Fig. 4), but transitions have increasingly affected cooler north-facing aspects across elevations (Fig. 5).

**Figure 5.**
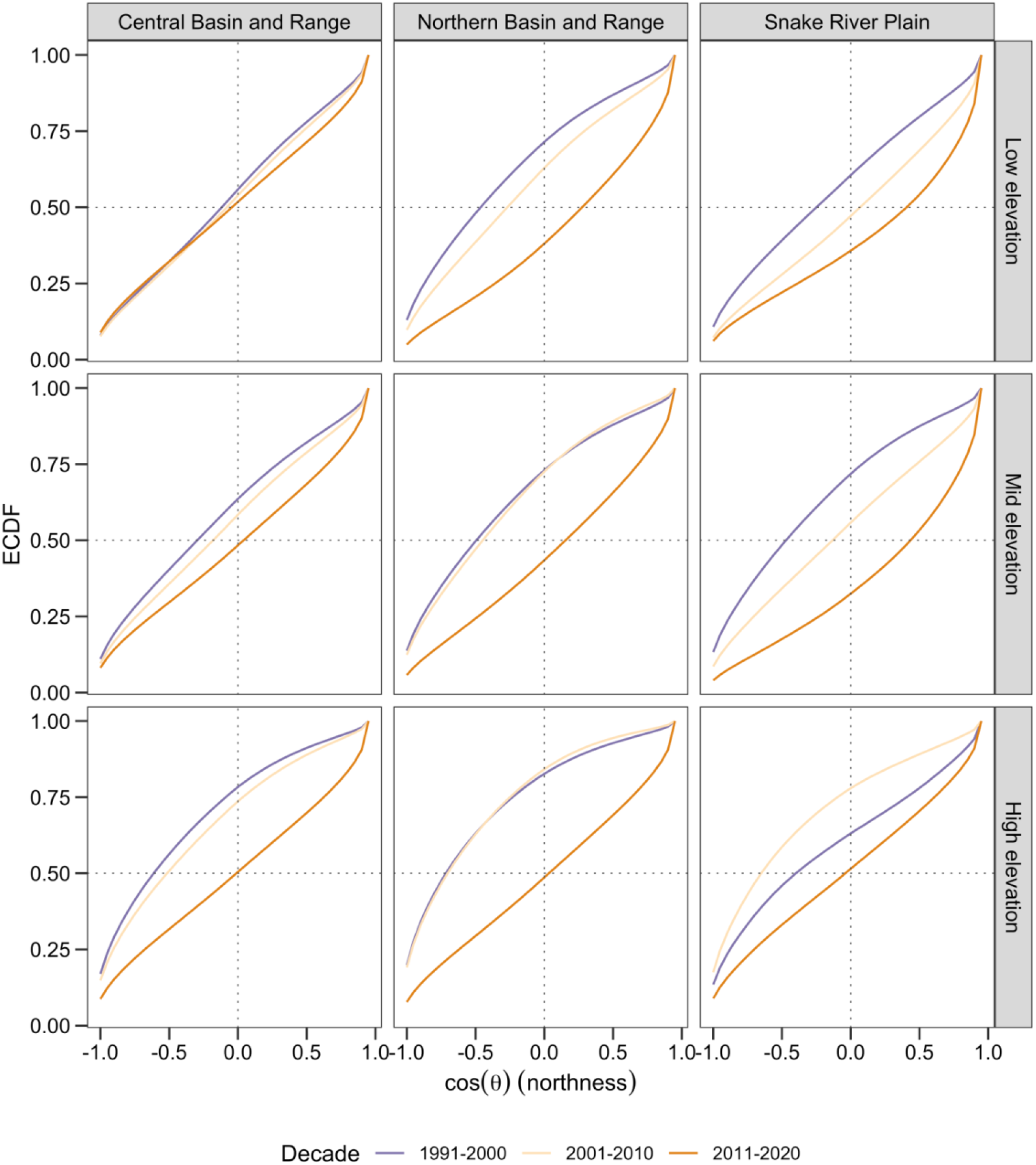
Distributional change in aspect of transitions to exotic annual grass dominance among elevational bands in three level III ecoregions of the Great Basin. Shifts toward more north-facing aspects (cos(θ) > 0) are evident across elevation bands and ecoregions.

**Figure 6.**
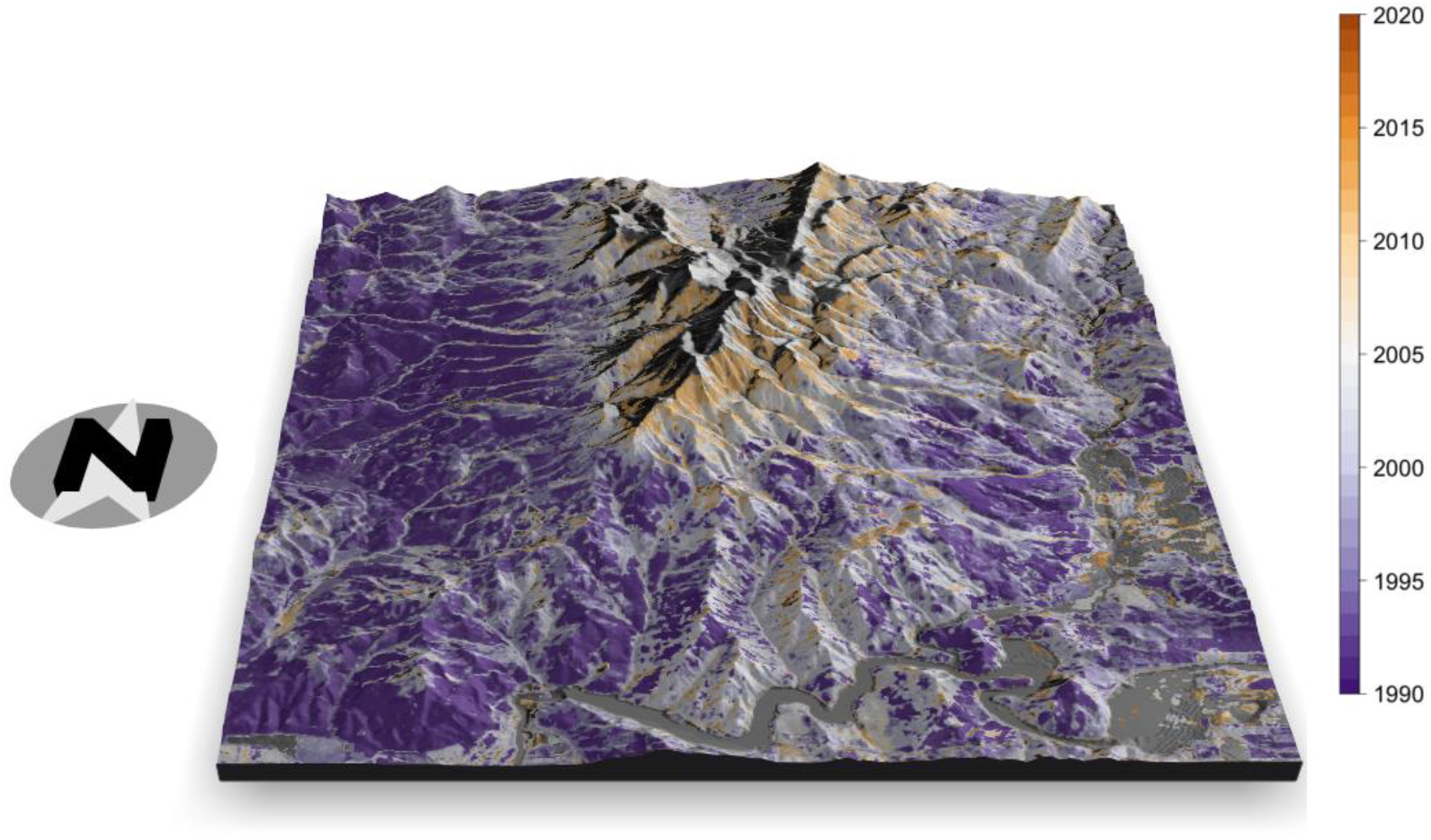
Example of elevational movement of annual grassland transitions north of Black Canyon Reservoir in Gem County, Idaho, USA. Non-annual grassland land cover and water in grey. At this site, the high elevation margin of annual grasslands ascended from approximately 1200 to 1700 m between 1990 and 2020.

Our analysis does not explicitly attribute these topographic shifts to a particular driver, but the patterns we observed are consistent with expected effects of warming based on considerable experimental research (Blumenthal et al., 2016; Compagnoni & Adler, 2014a, 2014b; Concilio et al., 2013; Zelikova et al., 2013). Upslope movement could also result from invasion proceeding outward from early-invaded areas, assuming initial invasion occurred at lower elevations. However, given that the earliest and most successful among the annual grass invaders, *B. tectorum*, achieved widespread distribution throughout the Great Basin by 1928 (Mack, 1981), and the rapid potential population growth rates of it and other annual grasses, it is doubtful that the upslope increase in dominance we observed is due only to a delayed legacy of initial introductions. Moreover, the interaction between aspect and elevation strongly suggests that environmental factors are at play: for instance, transitions to annual-dominance on north-facing slopes lags behind those on south-facing slopes by more than a decade across ecoregions (Fig. 4). Regardless of the mechanism(s), our findings suggest frequently invoked generalities about how susceptibility of rangelands to annual invasion changes across elevation gradients may not hold in the future (e.g., Chambers et al., 2007; Johnson et al., 2019).

The explosive expansion of annual grass dominance was unambiguous despite short-term fluctuations corresponding with increasingly severe episodes of spring drought (Fig. 3; Fig. S4). Consistent with the high interannual variability in cover of annuals previously described (Bradley & Mustard, 2005), short-term minima in extent observed in 1992, 2004, 2010, and 2015 predictably followed prolonged droughts (Fig. S4). Each retreat, however, was followed by an even higher maximum with the return of wet winters (Fig. 3). These short-term ebbs are probably most accurately understood as temporary expressions of low biomass and cover of adult plants, rather than transitions back to a previous state. Although droughts may temporarily inhibit their growth and reproduction, exotic annual grasses in the Great Basin have clearly benefited from the climate trajectory of the last 3 decades. Across the region, winters have warmed steadily with little change in precipitation, while summers have become drier (Fig. S4). In combination with periodic wet winters, warming is expected to benefit species such as *B. tectorum* (Compagnoni & Adler, 2014a), especially at higher elevations where temperature is limiting (Chambers et al., 2007), and these predictions are borne out at broad scales.

Among the most threatened biomes of North America (Knick et al., 2003), sagebrush and salt desert shrublands and their biota occupy a fragile elevational band in the Great Basin that is compressed between annual grassland transitions at lower elevations and woodland expansion predominantly at higher elevations (Miller et al., 2011). Our analysis reveals that this squeeze is tightening as annual grass dominance steadily moves upslope. Transformation of shrublands to exotic annual grasslands comes with dire socio-ecological consequences. Sagebrush and other shrubs lost in transitions are keystones supporting much of the region’s wildlife (Coates et al., 2016). Larger and more frequent wildfires directly threaten not only ecosystems, but also human health (Reisen et al., 2015; Wettstein et al., 2018). Both exotic annual grassland and woodland expansion threaten the forage base that sustains livestock operations and rural economies (Brunson & Tanaka, 2011).

The pace and scale of transitions supports the need for increased investment and strategic approaches to managing invasive annual grasses in the region (US Department of Agriculture, Natural Resources Conservation Service, 2020; Western Governors’ Association, 2020). Our findings suggest vulnerability to transition to annual grass dominance may be strongly controlled by broad-scale climatic drivers, highlighting the importance of this context— current and future—when considering alternative management actions. We recommend land managers prioritize efforts to proactively prevent less invaded rangelands from transitioning over reactive restoration of large-scale annual grasslands to their historical native plant communities, which is costly and ineffective (K. W. Davies et al., 2011; Pilliod et al., 2017). Such proactive management requires reducing exposure to annual grass seed sources (Sebastian et al., 2017), increasing resistance to invasion by promoting perennial plants (Chambers et al., 2014), and building adaptive capacity of local communities to respond early to the problem (Maestas et al., In Review). We acknowledge, however, that managers have few tools at their disposal and commonly lack sufficient resources to implement those tools at scales commensurate with the problem. Without increased investment and a paradigm shift in management mindset, the archetypal shrubland ecosystems of the Great Basin could be largely transformed into highly flammable, depauperate annual-dominated grasslands and woodlands within a lifetime.

## Acknowledgements

Funding for this research was provided by the US Department of Agriculture, Agricultural Research Service.

## Data accessibility statement

All data used in the analyses presented in this paper are freely available via the Rangeland Analysis Platform (https://rangelands.app/). Code used to process and extract the data via the Google Earth Engine code editor is provided in Supporting Information. The clustered time series is available to download at: http://rangeland.ntsg.umt.edu/data/rap/rap-derivatives/great-basin-classes/.

## Footnotes

2 Brazil’s Instituto Nacional de Pesquisas Espaciais (INPE) PRODES program annual deforestation estimates for the Brazillian Legal Amazon averaged 10,129 km^2^ from 2004–2019, or 0.2% of the 5 million km^2^ region. 2,373 km^2^ represents 0.6% of the total area of rangelands in our study area.

