## Supporting Information for "The elevational ascent and spread of invasive annual grass dominance in the Great Basin, USA"

**Table S1.** Candidate quantile regression models of elevations of transitions to annual grass dominance in the Great Basin, 1990–2020. Models for each quantile ($\tau$) are sorted in descending order of support according to AIC.

| $\tau$ | Model structure | log Likelihood | AIC |
|---|---|---|---|
| 0.1 | ecoregion + northness + year + ecoregion × northness × year | -25032.1 | 50088.1 |
| | ecoregion + northness + year + year × ecoregion | -25099.6 | 50213.3 |
| | ecoregion + northness + year + ecoregion × northness | -25436.6 | 50887.1 |
| | ecoregion + northness + year + year × northness | -25445.6 | 50903.1 |
| | ecoregion + northness + year | -25465.6 | 50941.1 |
| | ecoregion + northness + ecoregion × northness | -26351.4 | 52714.7 |
| | ecoregion + northness | -26371.3 | 52750.6 |
| 0.5 | ecoregion + northness + year + ecoregion × northness × year | -20989.6 | 42003.1 |
| | ecoregion + northness + year + year × ecoregion | -21072.4 | 42158.9 |
| | ecoregion + northness + year + year × northness | -21583.8 | 43179.6 |
| | ecoregion + northness + year + ecoregion × northness | -21671.6 | 43357.1 |
| | ecoregion + northness + year | -21679.7 | 43369.4 |
| | ecoregion + northness + ecoregion × northness | -23709.5 | 47430.9 |
| | ecoregion + northness | -23713.3 | 47434.6 |
| 0.9 | ecoregion + northness + year + ecoregion × northness × year | -31152.5 | 62329.0 |
| | ecoregion + northness + year + ecoregion × northness | -31200.5 | 62415.1 |
| | ecoregion + northness + year + year × ecoregion | -31801.3 | 63616.5 |
| | ecoregion + northness + year + year × northness | -31824.7 | 63661.5 |
| | ecoregion + northness + year | -31832.6 | 63675.2 |
| | ecoregion + northness + ecoregion × northness | -33431.6 | 66875.2 |
| | ecoregion + northness | -33911.8 | 67831.5 |

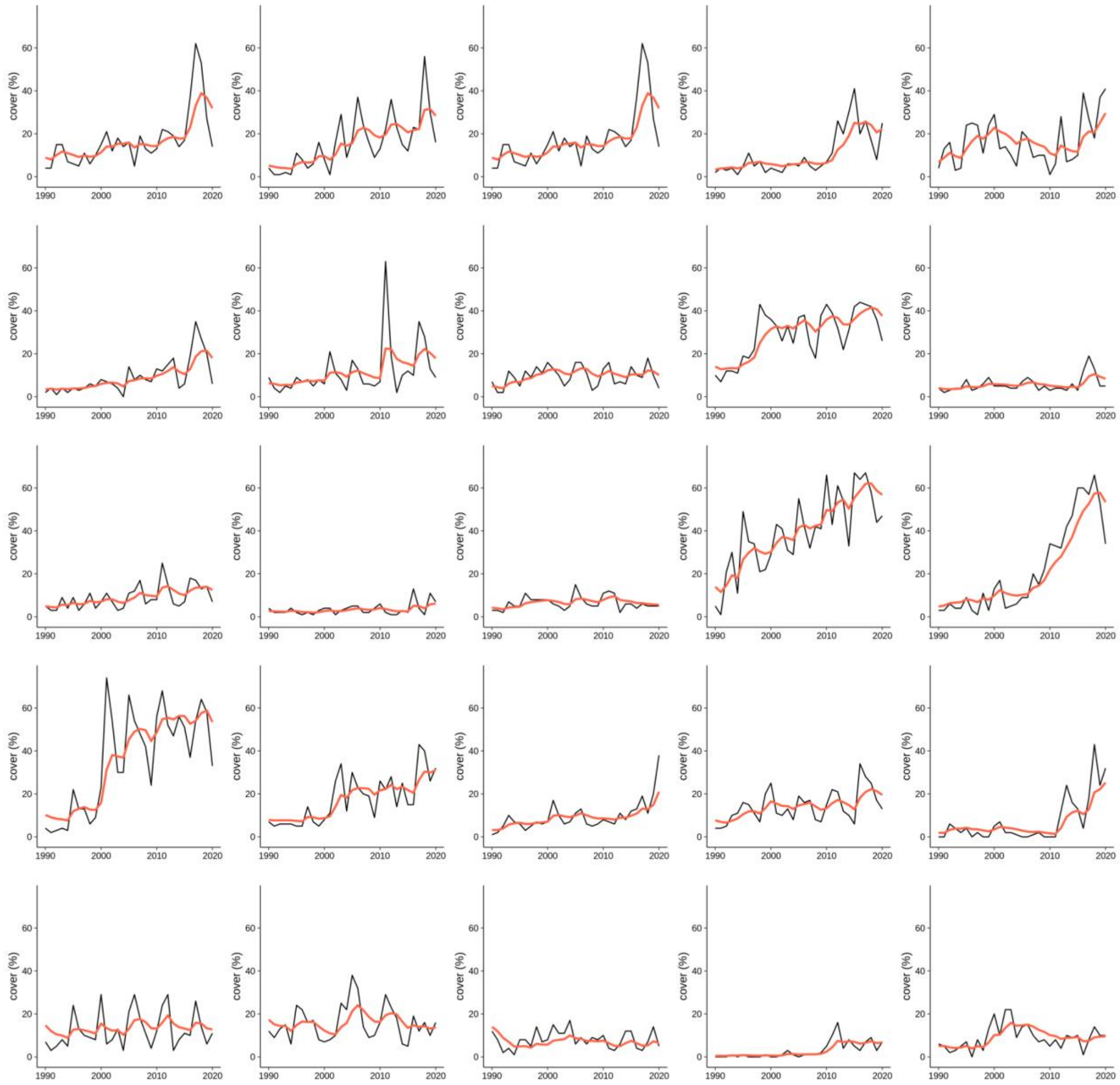


**Figure S1.** Smoothed (red) and raw (black) time series of annual forb and grass cover at 25 pixels selected manually to represent variation in annual forb and grass cover dynamics. Smoothing parameters that minimized mean absolute error among 1,000 randomly selected pixel time series were used to smooth the RAP cover estimates prior to unsupervised clustering. This dampened the magnitude of annual variation while still reflecting long-term trends and abrupt changes (e.g., due to wildfire or other disturbances).

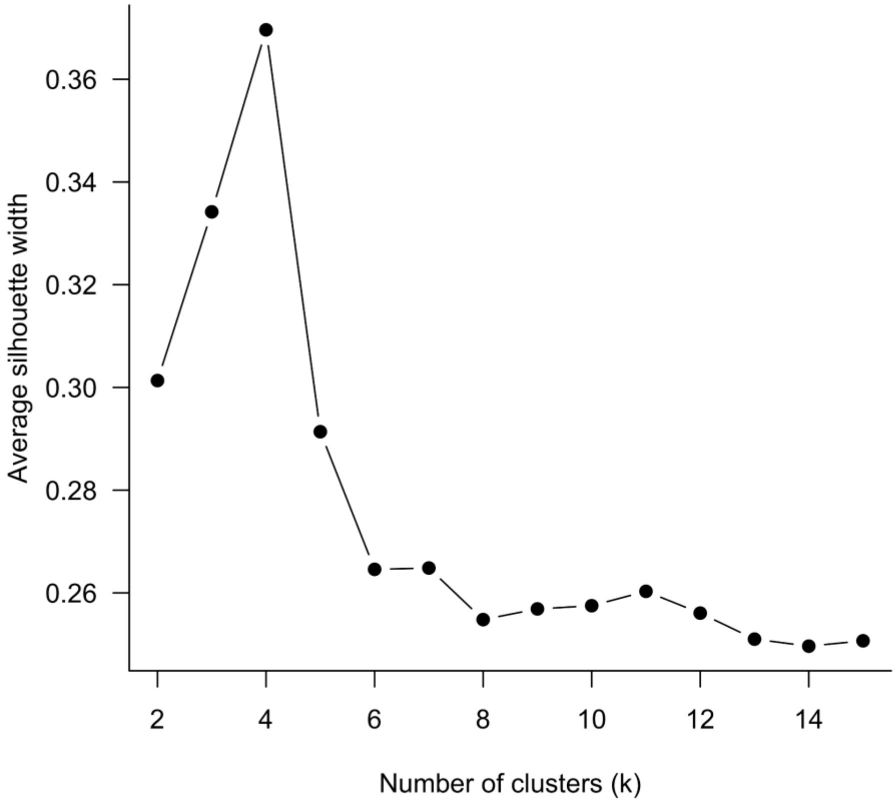


**Figure S2.** The silhouette method identified *k* = 4 as the most supported number of clusters to represent variation in functional group composition among rangelands in the Great Basin.

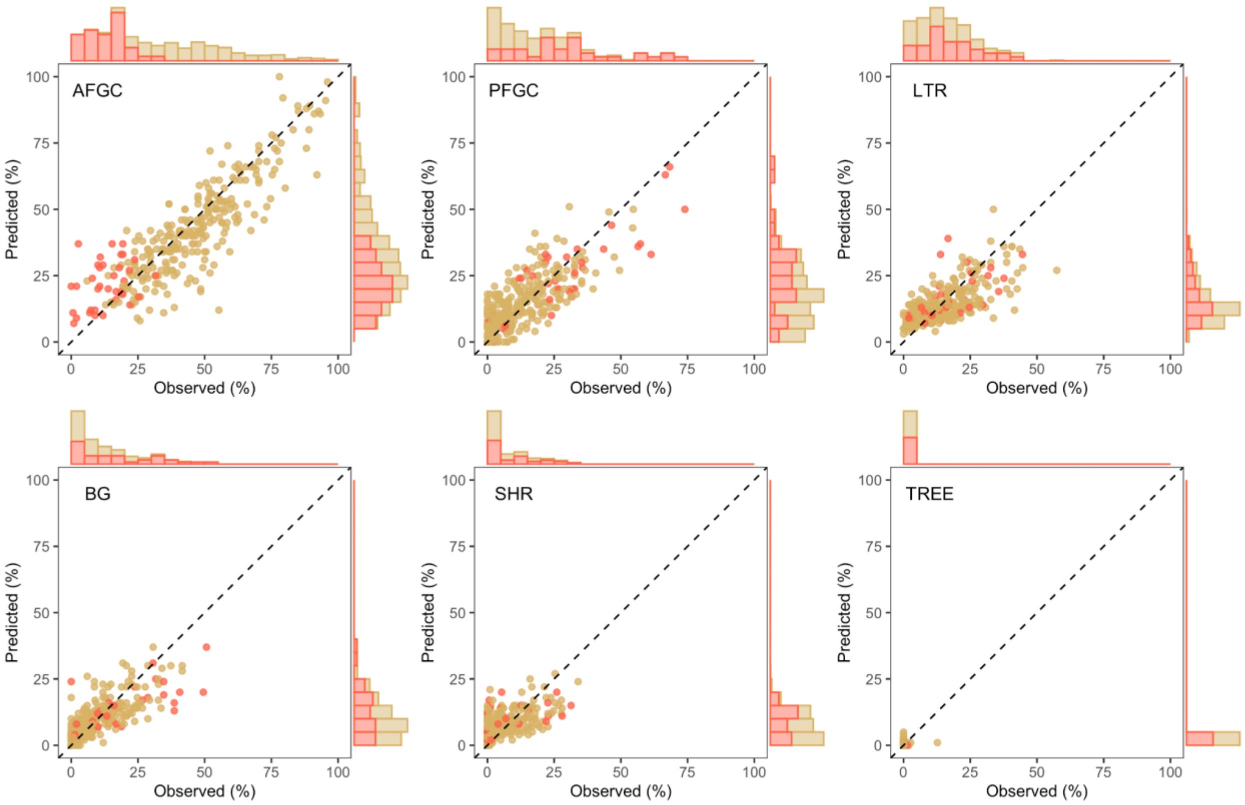


**Figure S3.** Field-estimated (x-axis) and RAP-estimated (y-axis) functional group cover among validation plots assigned to the annual grass dominance cluster based on RAP data. True positives (i.e., pixels also assigned to the annual grass dominance cluster based on field plot estimates; 88%) are shown in brown, false positives (i.e., pixels assigned to a non-annual grass dominance cluster based on field plot estimates; 12%) are shown in red. False positives were most often a result of overestimation of annual forb and grass cover by RAP relative to the field estimate.

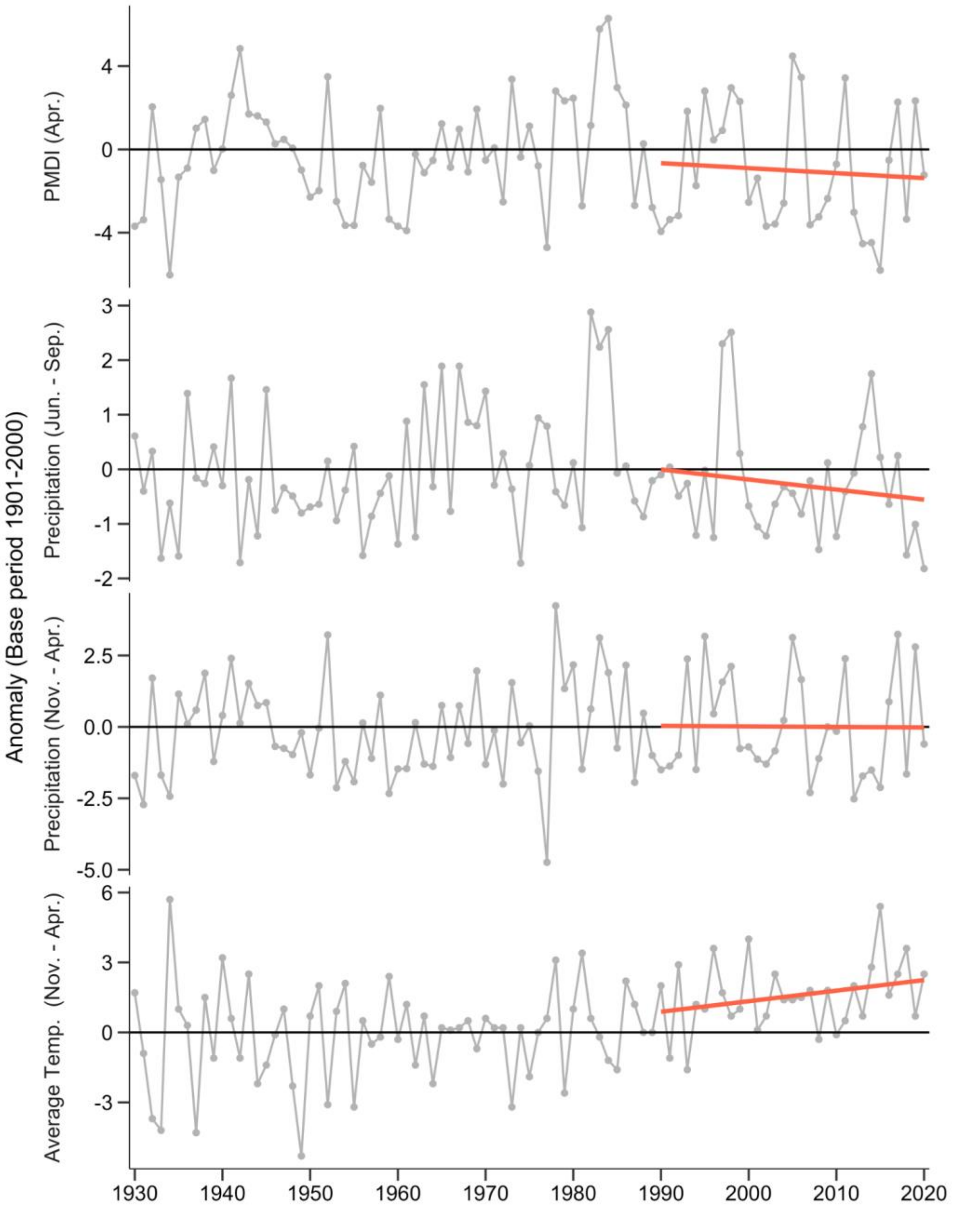


**Figure S4.** Trends in annual weather variables related to growth and reproduction of exotic annual grasses in the NOAA Great Basin region, 1930–2020. The decades over which we quantified expansion of annual grass dominance was characterized by periodic and increasingly severe drought events (PMDI = Palmer Modified Drought Index), decreasing summer precipitation, little change in average winter precipitation, and steadily increasing winter temperature. Red trend lines are least squares regressions fit to anomalies from 1990–2020.

**Appendix S1: Google Earth Engine code**

```
// Google Earth Engine code for analyses presented in the manuscript,
// "The elevational ascent and spread of invasive annual grass dominance
// in the Great Basin, USA"
//
// author: ***********
// contact: ***********
// revision date: August 11, 2021



// *** load assets, define study area and analysis mask *** //

// Load RAP v2 cover data.
var rapCover =
  ee.ImageCollection('projects/rangeland-analysis-platform/vegetation-cover-
v2')
    .select(['AFGC','PFGC','BG','SHR','LTR','TREE'])
    .filterDate('1984-01-01','2020-12-31');

// Store band names.
var coverBands = ee.List(['AFGC','PFGC','BG','SHR','LTR','TREE']);

// Study area:  Great Basin Ecoregions (level III).
var ecoregions = [
                  'Northern Basin and Range',
                  'Central Basin and Range',
                  'Snake River Plain'
                ];

// Load L3 ecoregion polygons.
var ecol3 = ee.FeatureCollection('EPA/Ecoregions/2013/L3');

// Select and union ecoregions into study area polygon.
var region = ecol3
  .filter(ee.Filter.inList('us_l3name',ecoregions))
  .union();

// Display study area.
Map.addLayer(ee.Image().paint(region, 0, 2), {}, 'region');

// Load USDA NASS Cropland Data Layer.
var cdl = ee.ImageCollection('USDA/NASS/CDL');

// Identify pixels classified as Grass/Pasture, Hay, or
// Alfalfa in >1 yr in USDA Cropland Data Layer (CDL).
var cdl_mask = cdl
  .select('cropland')
  .map(function(image){
    return image.eq(61)   // 61 = "Fallow/Idle Cropland"
    .or(image.eq(37))     // 37 = "Other Hay/Non Alfalfa"
    .or(image.eq(36));    // 36 = "Alfalfa"
  })
  .sum()
  .gt(1);  // must be in one of these classses in >1 yr
```

```
// Load rangeland land cover map based on Reeves, M. C., and J. E. Mitchell.
// 2011. Extent of coterminous US rangelands: Quantifying implications of
// differing agency perspectives. Rangeland Ecology & Management 64:585–597.
var rangeMap = ee.Image('projects/rangeland-analysis-platform/vegetation-
cover-mask-v1');

// Combine masks to limit analyses to rangeland landcover not in hay/alfalfa
class in CDL.
var rangeMask = rangeMap
          .lt(21)   // values >= 21 are forest, developed, or water (non-
rangeland)
          .and(cdl_mask.eq(0));
```

```
// ***  Step 1: smooth cover time series to emphasize long-term trends *** //

// Function to apply Holt's exponential smoothing method over an
ImageCollection.
var holtSmooth = function(thisImage, imageList) {

  // get the previous cover image from the end of the list
  var Img_tminus = ee.Image( ee.List(imageList).get(-1) );

  // names of all bands
  var coverBands = ee.List(['AFGC','PFGC','BG','SHR','LTR','TREE']);

  // names of bands used to store Beta (trend) parameters
  var bBands = ee.List(['B_AFGC','B_PFGC','B_BG','B_SHR','B_LTR','B_TREE']);

  // pull previous level l_tminus from last image
  var l_tminus = Img_tminus.select(coverBands);

  // pull previous trend b_tminus from last image
  var b_tminus = Img_tminus
    // select bBands
    .select(bBands)
    // rename them so they can be added to S bands later
    .rename(coverBands);

  // calculate new level l
  var l_t = thisImage
          .select(coverBands)
          .multiply(alpha) // current value * alpha
          .add(ee.Image(1)
                .subtract(alpha)
                .multiply(l_tminus.add(b_tminus))
              )           // + (1-alpha) * level @t-1
          // if smaller than zero, substitute zero (no negative cover)
          .max(ee.Image(0))
```

```
            .float();

  // calculate new trend b
  var b_t = l_t
            .subtract(l_tminus)
            .multiply(beta) // difference in level * beta
            .add(ee.Image(1)
                  .subtract(beta)
                  .multiply(b_tminus)
                )           // + (1-beta) * trend @t-1
            .float()
            // change band names back to bBands
            .rename(bBands);

  // combine level and trend bands into new img
  var Img_t = ee.Image(l_t)
    .addBands(b_t)
    .copyProperties(thisImage,['system:index','system:time_start']);

  return ee.List(imageList).add(Img_t);

};

// Make an image with initial trend (b) for each band in RAP cover.
// Initialize smoothing with b = 0.
var bBandsImage = ee.Image(0).select([0],['B_AFGC'])
    .addBands(ee.Image(0).select([0],['B_PFGC']))
    .addBands(ee.Image(0).select([0],['B_BG']))
    .addBands(ee.Image(0).select([0],['B_SHR']))
    .addBands(ee.Image(0).select([0],['B_LTR']))
    .addBands(ee.Image(0).select([0],['B_TREE']))
    .float();

// Set alpha (level smoothing parameter).
var alpha = 0.25;

// Set beta (trend smoothing parameter).
var beta = 0.01;

// Get first image in cover time series to initialize smoothing, put in a
list.
var rapFirst =  ee.List([
                  rapCover
                    // sort them chronologically
                    .sort('system:time_start')
                    // grab the first image (1984)
                    .first()
                    .select(coverBands)
                    // add bBands initialized with values of zero
                    .addBands(bBandsImage)
                ]);

// Get the rest of the time series to iterate over/
var rapSeries = rapCover
  .filterDate('1985-01-01','2020-12-31');
```

```
// Iterate Holt's smoothing function over time series.
var coverSmooth = ee.ImageCollection(
  ee.List(rapSeries.iterate(holtSmooth, rapFirst)))
  .select(coverBands)
  .map(function(image){ return image.float() });

Map.addLayer(coverSmooth.sort('system:index',true),{}, 'coverSmooth', false);




// ***  Step 2: sample training cases for clusterer  *** //

var firstYear =
ee.Date(rapCover.aggregate_min('system:time_start')).get('year');
var lastYear =
ee.Date(rapCover.aggregate_max('system:time_start')).get('year');
var trainYears = ee.List.sequence(firstYear, lastYear, 1);

// Get an image where each pixel is assigned a random
// index from 1 to length of trainYears.

// First, draw uniform random value in [0,1].
var tImage = ee.Image.random(0)
  // Then multiply by number of trainYears...
  .multiply(trainYears.size())
  // ...and round to nearest integer.
  .round()
  // Cast to integer data type.
  .toInt();

// Convert RAP imageCollection to an array image so we can slice out a random
// year at each pixel.
var rapArray = rapCover
  .sort('system:index')
  .select(coverBands)
  .toArray();

// Use random year index image tImage to slice out cover.
var tRap = rapArray
  .arraySlice( {
    // want to slice out a single row, so axis 0
    axis: 0,
    // want cover from row t
    start: tImage,
    // end is exclusive, so row t+1
    end: tImage.add(1)
  } )
  // arrayProject to convert from 2-D to 1-D array
  .arrayProject([1])
  // arrayFlatten to convert to image
  .arrayFlatten([coverBands]);
```

```
// ***  Step 3: build clusterer  *** //

// Select training cases, sampled randomly in time and space
// from raw RAP cover data (so covariance is among bands is captured).
var training = tRap
  .select(coverBands)
  // mask out non-rangelands
  .updateMask(rangeMask)
  // first oversample (some will be NA and dropped)
  .sample({
    region: region,
    scale: 30,
    numPixels: 3300,
    seed: 123,
    tileScale: 2,
    dropNulls: true,
    geometries: false
  })
  // then trim to desired size
  .limit(2500);

// Specify the clusterer and train it.
var clusterer = ee.Clusterer.wekaKMeans({
  nClusters: 4,
  init: 1,
  distanceFunction: 'Euclidean',
  maxIterations: 10
}).train(training);




// ***  Step 4: apply clusterer to smoothed time series  *** //

var clusterCollection = coverSmooth
  .filterDate('1990-01-01','2020-12-31') // first 5 years used to initialize
smoothing
  .map(function(image){
  return image
    .cluster(clusterer)
    .copyProperties(image, ['system:time_start','system:index']);
});


Map.addLayer(clusterCollection, {min:0, max:3}, 'clusters', false);




// ***  Step 5: convert cluster IDs to binary annual grass dominance/other
*** //

// Specify which cluster label represents annuals-dominated state (cluster
0).
```

```
var anualDomCluster = ee.Number(0);

// Map over clustered time series, converting to binary AGL
var annualDom = clusterCollection.map(function(image){
  var year = image.date().get('year').toString();
  return image
    .eq(anualDomCluster)
    .select([0],['ad'])
    .copyProperties(image, ['system:time_start'])
    .set('system:index', year);
});




// ***  Step 6: identify first transition year  *** //

var transition = annualDom
  // map a function over the collection that replaces 1s with the year of the
image.
  .map(function(image){
    var thisYear = image.date().get('year');
    return image
            .remap({
              from:[1],
              to:[ee.Number(thisYear)],
              defaultValue:9999,
              bandName:'ad'
            });
  })
  // min finds first year (i.e., year of 1st transition)
  .min()
  .select([0],['ad1']) // rename band 'ad1'
  // add a mask band to show only pixels appearing in annual dominance
cluster
  // in at least 1 year
  .addBands(annualDom.sum().gte(1).select([0],['datamask']));


// Define some visualization parameters for this image.
var visParams = {
    min:1990,
    max:2020,
    palette:
['542788','8073ac','b2abd2','d8daeb','f7f7f7','fee0b6','fdb863','e08214','b35
806']
  };

// Add to map.
Map.addLayer(transition.select('ad1')
  .updateMask(transition.select('datamask'))
  .updateMask(rangeMask)
  .clip(region),
  visParams,
  'First Year Transitioned', true);
```

```
// ***  Step 7: sample elevation/aspect stratified by transition year image
*** //

// Add 'code' property (numeric) to ecoregion FeatureCollection.
ecol3 = ecol3.map(function(feature) {
  var code = feature.get('us_l3code');
  var numCode = ee.Number.parse( code ).byte();
  return feature.set('code',numCode);
});

// Convert ecoregion FeatureCollection to an image, taking 'code' value.
var regionImage = ecol3
  .filter(ee.Filter.inList('us_l3name',ecoregions))
  .reduceToImage({
    properties: ['code'],
    reducer: ee.Reducer.first(),
  })
  .toInt()
  .select([0],['region']);

// Get elevation image (USGS National Elevation Dataset 1/3 arc-second).
var elevation = ee.Image('USGS/NED').round().toUint16();

// Calculate aspect from DEM.
var aspect = ee.Terrain.aspect(elevation).float();

// Sample elevations and aspect by transition year to analyze in R.
var sampleTransitionTopo = elevation
      .round()
      .toInt()
      .addBands(aspect)
      .addBands(regionImage)
      .addBands(transition.select('ad1'))
      .updateMask(rangeMask)
      .stratifiedSample({
        numPoints: 5e3, // way oversample to get large enough sample in each
ecoregion.
        classBand: 'ad1',
        region: region.geometry(),
        scale: 30,
        seed: 123,
        tileScale: 4,
        geometries: false
      });

// Export.
Export.table.toDrive({
  collection: sampleTransitionTopo,
  description: 'sampleTransitionTopo',
  fileFormat: 'CSV'
});
```

```
// ***  Step 8: calculate annual grassland area in each year  *** //

// Map over annualDom collection, calculating area in annual dominance and
// storing in a geometry-less feature for each year.
var annualDom_area = ee.FeatureCollection(
  annualDom.map(function(img){

    // get year.
    var year = img.get('system:index');

    // multiply binary annualDom image by pixelArea.
    var area = ee.Image.pixelArea().multiply(img);

    // reduce over regions.
    var reduced = area
      .updateMask(rangeMask)
      .reduceRegions({
        collection: ecol3.filter(ee.Filter.inList('us_l3name',ecoregions)),
        reducer: ee.Reducer.sum(),
        scale: 30,
        tileScale: 4
      });

    // add a 'year' column and strip geometries.
    reduced = reduced
      .map(function(feature) { return feature.set('year',year) })
      .select(['.*'],null,false);

    return reduced;

  })
  ).flatten();


// Export.
Export.table.toDrive({
  collection: annualDom_area,
  description: 'annualDom_area',
  fileFormat: 'CSV'
});




// *** Step 9: Get ECDFs of aspects of transitions for each decade of time
series *** //

// Convert aspect to 'northness'
var northness = aspect
  .multiply(3.14159)
  .divide(180)
  .cos()
  .rename('northness');

// Calculate slope (mask out flat terrain);
```

```
var slope = ee.Terrain.slope(elevation);

// function to get histogram data
var getHistData = function(image, band, roi, min, max, steps) {

  var histogram = image
    .select(band)
    .reduceRegion({
      reducer: ee.Reducer.fixedHistogram({
        min: min,
        max: max,
        steps: steps,
        cumulative: false
      }),
      geometry: roi,
      scale: 30,
      maxPixels: 1e9,
      tileScale: 2
  });

  // Get the list of DN values (x-axis of the histogram)
  var dnList = ee.Array(histogram.get(band)).slice(1,0,1);
  // Get the list of Counts values (y-Axis of the histogram)
  var Counts = ee.Array(histogram.get(band)).slice(1,1);
  // Calculate sum of counts.
  var totalCount = Counts.reduce({reducer: ee.Reducer.sum(), axes:
[0]}).project([0]);
  // Divide each value by the total so that the values are between 0 and 1
  // This will be the probability at each DN
  var Proportions = ee.Array(Counts).divide(totalCount.get([0]));

  // Create a merged array with DN and probabilities
  var array = ee.Array.cat({arrays: [ dnList,
                                      Counts,
                                      Proportions
                                    ],
                            axis:1
                            });

  // Convert the array into a feature collection with null geometries
  var fc = ee.FeatureCollection(array.toList().map(function(list) {
    return ee.Feature(null, {
      dn: ee.List(list).get(0),
      count: ee.List(list).get(1),
      proportion: ee.List(list).get(2)
    });
  }));

  return fc;

};

// map over ecoregions
var ecos = ecol3
  .filter(ee.Filter.inList('na_l3name', ecoregions));

var eco_col = ecos
```

```
.map(function(feature){

  var econame = feature.get('na_l3name');

  // for a given ecoregion, compute tercile breaks of elevations
  // of all annual-dominated pixels across time.
  var breaks = elevation
    .updateMask(annualDom.sum().gte(1))
    .reduceRegion({
      reducer: ee.Reducer.percentile([33,66]),
      geometry: feature.geometry(),
      scale: 240,
      maxPixels: 1e9,
      tileScale: 2
    })
    .toImage();

  // convert to categorical image
  var elevation_bands = ee.Image(0)
    .where(elevation
            .gt(breaks.select(0))
            .and(elevation.lte(breaks.select(1))),
            ee.Image(1))
   .where(elevation
            .gt(breaks.select(1)),
            ee.Image(2));

  var band_col = ee.FeatureCollection(ee.List([0,1,2])
    .map(function(band){

      // for a given band, calculate histograms for early (1991-2000)
      // and late (2011-2020) periods
      var earlyMask = transition.select('ad1').gt(1990)
        .and(transition.select('ad1').lte(2000))
        .and(slope.gt(5))
        .and(elevation_bands.eq(ee.Image.constant(band)));

      var earlyHist = getHistData(
        northness.updateMask(earlyMask).float(),
        'northness', feature.geometry(),
          -1,1,40);

      var middleMask = transition.select('ad1').gt(2000)
        .and(transition.select('ad1').lte(2010))
        .and(slope.gt(5))
        .and(elevation_bands.eq(ee.Image.constant(band)));

      var middleHist = getHistData(
        northness.updateMask(middleMask).float(),
        'northness', feature.geometry(),
          -1,1,40);

      var lateMask = transition.select('ad1').gt(2010)
        .and(transition.select('ad1').lte(2020))
        .and(slope.gt(5))
        .and(elevation_bands.eq(ee.Image.constant(band)));
```

```
        var lateHist = getHistData(
          northness.updateMask(lateMask).float(),
          'northness', feature.geometry(),
            -1,1,40);

        // pull relevant columns
        var northnessList = ee.Array(earlyHist.aggregate_array('dn'))

        var earlyList = ee.Array(earlyHist.aggregate_array('proportion'));

        var middleList = ee.Array(middleHist.aggregate_array('proportion'));

        var lateList = ee.Array(lateHist.aggregate_array('proportion'));

        // Create a merged array with data
        var array = ee.Array.cat({arrays: [
                                    northnessList,
                                    earlyList,
                                    middleList,
                                    lateList
                                  ],
                             axis: 1
                           });

        // Convert the array into a feature collection with null geometries
        var fc = ee.FeatureCollection(array.toList().map(function(list) {
          return ee.Feature(null, {
            elevation_band: band,
            ecoregion: econame,
            northness: ee.List(list).get(0),
            earlyProportion: ee.List(list).get(1),
            middleProportion: ee.List(list).get(2),
            lateProportion: ee.List(list).get(3)
          });
        }));

        return fc;

      })).flatten();  // flatten over elevation bands

    return band_col;

  }).flatten();  // flatten over ecoregions

// Export.
Export.table.toDrive({
  collection: eco_col,
  description: 'northness_by_elevation_band',
  fileFormat: 'CSV'
});
```
